# The nucleus accumbens core is necessary to scale fear to degree of threat

**DOI:** 10.1101/2020.02.06.917328

**Authors:** Madelyn H. Ray, Alyssa N. Russ, Rachel A. Walker, Michael A. McDannald

## Abstract

Fear is adaptive when the level of the response rapidly scales to degree of threat. Using a discrimination procedure consisting of danger, uncertainty and safety cues, we have found rapid fear scaling (within two seconds of cue presentation) in male rats. Here we examined a possible role for the nucleus accumbens core (NAcc) in the acquisition and expression of fear scaling. In experiment 1, male Long Evans rats received bilateral sham or neurotoxic NAcc lesions, recovered and underwent fear discrimination. NAcc-lesioned rats were generally impaired in scaling fear to degree of threat, and specifically impaired in rapid uncertainty-safety discrimination. In experiment 2, male Long Evans rats received NAcc transduction with halorhodopsin or a control fluorophore. After fear scaling was established, the NAcc was illuminated during cue or control periods. NAcc-halorhodopsin rats receiving cue illumination were specifically impaired in rapid uncertainty-safety discrimination. The results reveal a general role for the NAcc in scaling fear to degree of threat, and a specific role in rapid discrimination of uncertain threat and safety.

**Significance Statement:** Rapidly discriminating cues for threat and safety is essential for survival and impaired threat-safety discrimination is a hallmark of stress and anxiety disorders. In two experiments, we induced nucleus accumbens core (NAcc) dysfunction in rats receiving fear discrimination consisting of cues for danger, uncertainty and safety. Permanent NAcc dysfunction, via neurotoxic lesion, generally disrupted the ability to scale fear to degree of threat, and specifically impaired one component of scaling: rapid discrimination of uncertain threat and safety. Reversible NAcc dysfunction, via optogenetic inhibition, specifically impaired rapid discrimination of uncertain threat and safety. The results reveal that the NAcc is essential to scale fear to degree of threat, and is a plausible source of dysfunction in stress and anxiety disorders.

## Introduction

The ability to discriminate danger from safety is critical to survival. Individuals with stress and anxiety disorders are impaired in discrimination, showing excessive fear-related responses to safety (Jovanovic et al., 2010; Jovanovic et al., 2012; Lissek et al., 2014; Duits et al., 2015). Danger and safety represent extremes of a threat continuum, with most real-world threats involving uncertainty. Ideally, one’s level of fear should scale to the degree of threat. A scaled fear response would be most adaptive if it was rapidly organized following encounter with a potential threat.

Drawing from learning theory (Rescorla, 1968), our laboratory devised a discrimination procedure in which distinct auditory cues predict unique foot shock probabilities: danger (p=1.00), uncertainty (p=0.25), and safety (p=0.00) (Berg et al., 2014). Using this procedure, we have found that fear level scales to shock probability within two seconds of cue presentation (DiLeo et al., 2016). Present work in our laboratory seeks to identify brain regions necessary for fear scaling and its rapid emergence. Candidate regions should be able to process valence and receive amygdalar input (Quirk et al., 1995; Goosens and Maren, 2001; Koo et al., 2004; McDannald and Galarce, 2011). We identified the nucleus accumbens core (NAcc) as a likely candidate, based on its ability to rapidly process reward-predictive cues (Cromwell and Schultz, 2003; Setlow et al., 2003; Ambroggi et al., 2011; McGinty et al., 2013; Saddoris and Carelli, 2014; Sugam et al., 2014; Ottenheimer et al., 2018), as well as its anatomical connectivity with the amygdala (Kita and Kitai, 1990; Petrovich et al., 1996; Wright and Groenewegen, 1996). Even more, the NAcc is implicated in a variety of fear-related processes (Haralambous and Westbrook, 1999; Thomas et al., 2002; Schwienbacher et al., 2004; Iordanova et al., 2006b; Fadok et al., 2010; Badrinarayan et al., 2012; Oleson et al., 2012; Li and McNally, 2015; Correia et al., 2016).

In two experiments, we examined roles for the NAcc in fear scaling. In experiment 1, we permanently ablated NAcc neurons via neurotoxic lesion. Following recovery, rats received fear discrimination consisting of danger, uncertainty, and safety cues. Fear was measured with suppression of rewarded nose poking (Estes and Skinner, 1941; Bouton and Bolles, 1980). Examining suppression over the entire 10-s cue permitted analysis of overall fear scaling. To examine the temporal emergence of scaling, we divided the 10-s cues into five, 2-s cue intervals. Focusing on suppression during the first 2-s cue interval permitted analysis of rapid fear scaling. In experiment 2, we examined a role for the NAcc in the expression of fear scaling. Rats were NAcc-transducted with halorhodopsin or a control fluorophore, and bilaterally implanted with ferrules above the NAcc. Following recovery, rats received fear discrimination until fear scaling was stable. Over the next eight sessions, the NAcc was green-light illuminated during cue presentation or a control period, optogenetically inhibiting activity in halorhodopsin rats. The two experiments allowed us to examine roles for the NAcc in the acquisition and expression of fear scaling.

## Materials and methods

### Experiment 1

#### Subjects

Subjects were forty-five male Long Evans rats weighing 275-300 g upon arrival (Charles River Laboratories; RGD Cat# 2308852, RRID:RGD_2308852). Rats were individually housed and maintained on a 12-h dark light cycle (lights off at 6:00 PM) with water *ad libitum*. Procedures adhered to the NIH Guide for the Care and Use of Laboratory Animals and were approved by the Boston College Institutional Animal Care and Use Committee.

#### Behavioral apparatus

Eight sound-attenuated enclosures each housed a behavior chamber with aluminum front and back walls, clear acrylic sides and top, and a metal grid floor. Grid floors were electrically connected to a shock generator. A single external food cup and central nose poke opening equipped with infrared photocells were present on one wall. Auditory stimuli were presented through two speakers mounted on the ceiling of each behavior chamber.

#### Surgical procedures

Stereotaxic surgery was performed under isoflurane anesthesia (2-5%) using aseptic technique. Twenty-four rats received bilateral infusions of *N*-Methyl-D-aspartic acid (15 μg/μl in Dulbecco’s PBS) aimed at the nucleus accumbens core (0.40 μl, +1.90 AP, ±1.80 ML, −6.60 DV from skull). Infusions were delivered via 2 μl syringe (Hamilton, Neuros) controlled by a microsyringe pump (World Precision Instruments, UMP3-2). Infusion rate was ~0.11 μl/min. Thirty seconds after the completion of each infusion, the syringe was raised 0.1 mm then left in place for five minutes to encourage delivery to the target site. The remaining twenty-one rats received identical surgical treatment without infusions. Rats received carprofen (5 mg/kg) for post-operative analgesia.

#### Nose poke acquisition

Following recovery from surgery, rats were food restricted to 85% of their initial free feeding body weight, then fed (2 - 20 g/day) to increase their target body weight by 1 g/day for the remainder of testing. Rats were shaped to nose poke for pellet (BioServ F0021 – protein/fat/carbohydrate blend) delivery using a fixed ratio 1 schedule: one nose poke yielded one pellet. Shaping sessions lasted 30 min or approximately 50 nose pokes. Over the next 3, 60-min sessions, rats were placed on variable interval (VI) schedules in which nose pokes were reinforced on average every 30 s (session 1), or 60 s (sessions 2 and 3). For the remainder of testing, nose pokes were reinforced on a VI-60 schedule independent of all Pavlovian contingencies.

#### Pre-exposure

In two separate sessions, each rat was pre-exposed to the three cues to be used in Pavlovian fear discrimination. Cues were auditory stimuli, 10-s in duration and consisted of repeating motifs of a broadband click, phaser, or trumpet. Stimuli can be heard or downloaded at http://mcdannaldlab.org/resources/ardbark. Previous studies have found these stimuli to be equally salient, yet highly discriminable (Berg et al., 2014; Wright et al., 2015; DiLeo et al., 2016; Ray et al., 2018). The 42-min pre-exposure sessions consisted of four presentations of each cue (12 total presentations) with a mean inter-trial interval (ITI) of 3.5 min. The order of trial type presentation was randomly determined by the behavioral program and differed for each rat during each session throughout behavioral testing.

For all sessions, fear to each auditory cue was measured using a suppression ratio based on nose poke rates during the 20-s baseline period immediately preceding the 10-s cue period: suppression ratio = (baseline nose poke rate – cue nose poke rate) / (baseline nose poke rate + cue nose poke rate). A ratio of 1 indicated complete suppression of nose poking during the cue and a high level of fear; 0, no suppression and no fear. Intermediate suppression ratios reflected intermediate fear levels. The same suppression ratio formula was used to calculate fear in 2-s cue intervals.

#### Fear discrimination

Each rat received sixteen, 54-min Pavlovian fear discrimination sessions. Sessions began with a ~5-min warm-up period during which no cues or shock were presented. The three cues were associated with a unique foot shock (0.5 mA, 0.5-s) probability: danger (1.00), uncertainty (0.25), and safety (0.00). Foot shock was administered 1-s following cue offset. A single session consisted of four danger, six uncertainty omission, two uncertainty shock, and four safety trials. Auditory stimulus identity was counterbalanced across rats. Mean inter-trial interval was 3.5 min.

#### Histology

Upon the conclusion of behavior, rats were anesthetized with an overdose of isoflurane and perfused intracardially with 0.9% biological saline. Brains were extracted and stored in 4% (v/v) formalin and 10% (w/v) sucrose. Forty-micrometer sections were collected on a sliding microtome. Tissue was then washed with PBS, incubated in NeuroTrace (Thermo Fisher, N21479) at a 1:200 concentration, washed again, mounted, dried, and coverslipped with Vectashield Hardset mounting media (Vector Labs, H-1400). Slides were imaged within 3 weeks of processing.

### Experiment 2

#### Subjects

Subjects were 25 male Long Evans rats weighing 275-300 g upon arrival (Charles River Laboratories; RGD Cat# 2308852, RRID:RGD_2308852). Rats were individually housed and maintained on a 12-h dark light cycle (lights off at 6:00 PM) with water *ad libitum*. Procedures adhered to the NIH Guide for the Care and Use of Laboratory Animals and were approved by the Boston College Institutional Animal Care and Use Committee.

#### Behavioral apparatus

Behavioral apparatus was identical to experiment 1. In addition to the standard behavior apparatus, green lasers (532 nm, max 500 mW; Shanghai Laser & Optics Century Co., Ltd.; Shanghai, China) were used to illuminate the NAcc. Lasers were connected to the behavior cables via 1X2 fiber optic rotatory joints (Doric; Quebec, Canada). A ceramic sleeve maintained contact between the ferrules on the optogenetic cable and the head cap. The ferrule junction was shielded with black shrink wrap to block light emission into the behavioral chamber. A PM160 light meter (Thorlabs; Newton, NJ) was used to measure light output.

#### Optogenetic materials

Optical ferrules were constructed using 2.5mm ceramic zirconia ferrules (Precision Fiber Products; Chula Vista, CA). Behavior cables were custom made for light delivery (Multimode Fiber, 0.22 NA, High-OH, Ø200 μm Core). All protocols can be downloaded at http://mcdannaldlab.org/resources/optogenetics.

#### Surgical procedures

Stereotaxic surgery was performed under isoflurane anesthesia (2-5%) using aseptic technique. Thirteen rats received bilateral infusions of AAV-hSyn-eNpHR3.0-EYFP (*halorhodopsin*) aimed at the nucleus accumbens core (0.50 μl, +1.90 AP, ±1.80 ML, −6.60 DV at a 0° angle) and bilateral optical ferrules (+1.70 AP, ±2.80 ML, −6.00 DV at a 10° angle). Infusions were delivered via 2 μl syringe (Hamilton, Neuros) controlled by a microsyringe pump (World Precision Instruments, UMP3-2). Infusion rate was ~0.11 μl/min. The syringe was raised 0.1 mm after each infusion, then left in place for five min to encourage delivery to the target site. The remaining 12 rats received identical surgical treatment but were infused with a control fluorophore (AAV-hSyn-EYFP). Implants were secured with dental cement surrounded by a modified, 50 mL centrifuge tube. Post-surgery, rats received 2 weeks of undisturbed recovery with prophylactic antibiotic treatment (cephalexin; Henry Schein 049167) before beginning nose poke acquisition. All rats received carprofen (5 mg/kg) for post-operative analgesia.

#### Pre-illumination training and cable habituation

Nose poke acquisition, pre-exposure and initial fear discrimination (10 sessions) were identical to Experiment 1. We increased the delay between cue offset and shock onset to 2 s to ensure that neural activity would not be inhibited during shock delivery. Cable habituation was provided in two consecutive sessions by plugging rats into optogenetic cables and administering fear discrimination without illumination. In total, rats received twelve fear discrimination sessions prior to receiving light illumination.

#### NAcc illumination

Rats received eight sessions of fear discrimination plus NAcc illumination. The NAcc was illuminated via bilateral delivery of 12.5 mW of 532 nm ‘green’ light: DPSS laser → optogenetic cables → implanted ferrules. There were two types of illumination sessions: cue and ITI. For cue sessions, light illumination began 0.5 s prior to cue onset and ended 0.5 s following cue offset, resulting in a total illumination time of 11 s. Light illumination was given for all trial types (danger, uncertainty and safety) for a total of 16 illumination events per session. For ITI sessions, illumination occurred during the inter-trial intervals between cue presentations. Illumination was roughly equidistant from previous cue offset and subsequent cue onset (~90 s from each). Sixteen ITI illumination events were administered, each lasting 11 s, equating total illumination time for cue and ITI sessions. The within-subjects design meant that each rat received four cue illumination sessions and four ITI illumination sessions. Illumination was given in two-session blocks, with half of the subjects starting with cue illumination.

#### Histology

After behavioral testing ended, rats were anesthetized with an overdose of isoflurane and perfused intracardially with 0.9% biological saline and 4% paraformaldehyde in a 0.2 M potassium phosphate buffered solution. Brains were extracted and stored in 4% (v/v) formalin and 10% (w/v) sucrose. Forty-micrometer sections were collected on a sliding microtome. Tissue was rinsed, incubated in NeuroTrace (Thermo Fisher, N21479) at a 1:200 concentration, rinsed again, mounted, dried, and coverslipped with Vectashield Hardset (Vector Labs, H-1400). Slides were imaged within 3 weeks of processing.

#### Statistical analysis

Behavioral data were acquired using Med Associates Med-PC IV software (MED PC, RRID:SCR_012156). Raw data were processed in Matlab (MATLAB, RRID:SCR_001622) to extract time stamps for nose poke and cue onset. Suppression ratios were calculated as: (baseline poke rate – cue poke rate) / (baseline poke rake + cue poke rate) and were analyzed with repeated measures ANOVA in SPSS (RRID:SCR_002865). Repeated measures ANOVA was performed with factors of group, cue, and time (experiment 1) or group, cue, time, and illumination (experiment 2). Partial eta squared (η_p_^2^) and observed power (op) are reported for ANOVA results for indicators of effect size. For all analyses, *p*<0.05 (or an appropriate Bonferroni correction) was considered significant.

## Results

### Experiment 1

#### Histological Results

Rats received bilateral sham or neurotoxic NAcc lesions. Neurotoxic damage (cell loss and gliosis) was quantified. Twenty-four NAcc rats showed damage primarily in the NAcc (>85%) with minor damage (~10% or less) in the neighboring accumbens shell. Shams showed no evidence of neurotoxic damage. Representative sham (Figure 1A, left), and NAcc lesion (Figure 1A, right) sections are shown. Each subject’s lesion was drawn, made transparent, and stacked (Figure 1B). Darker areas indicate regions of greater overlap and more consistent damage. Rats fully recovered from surgery before receiving fear discrimination (Figure 1C).

**Figure 1.**
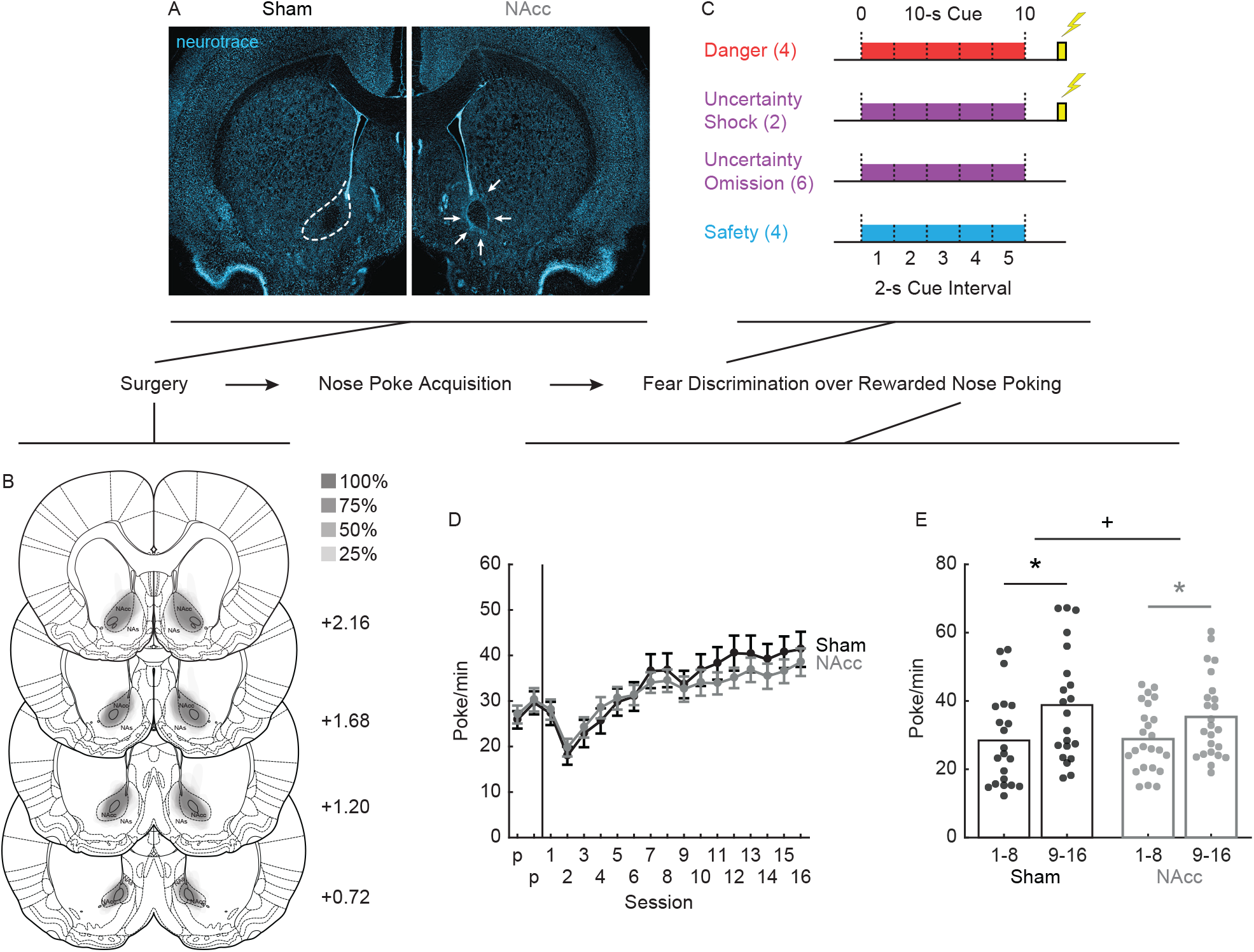
NAcc lesion experimental outline. (A) Representative sham with NAcc intact (left) and lesion with NAcc damage (right) is shown. Dotted lines (left) show approximate NAcc location. Arrows (right) indicate gliosis and damage restricted to the NAcc. (B) The extent of neurotoxic NAcc lesions across four coronal planes is shown, and the anterior distance from bregma (millimeters) indicated. (C) Pavlovian fear discrimination consisted of three, 10-s cues predicting unique foot shock probabilities: danger (p=1.00), red; uncertainty (p=0.25), purple; and safety (p=0.00), blue. Cues were divided into 5, 2-s intervals for rapid analyses. (D) Mean ± SEM baseline nose poke rates for the sixteen fear discrimination sessions are shown for sham (black) and NAcc (gray) rats. (E) Mean baseline nose poke rates for sessions 1-8 and 9-16 for sham and NAcc rats. Data points show individual poke rates. *independent samples t-test, p<0.025, +block × group interaction p<0.05. Abbreviations: NAcc – nucleus accumbens core, NAs – nucleus accumbens shell.

#### Baseline Nose Poking

NAcc lesions altered the progression of nose poking over discrimination sessions, but did not grossly reduce nose poke rates (Figure 1D). ANOVA for baseline nose poke rate with session (16) and group (sham vs. NAcc) as factors found a main effect of session (F_(15,645)_ = 47.14, *p*=3.77 × 10^−93^, η_p_^2^ = 0.52, op = 1.00), a session × group interaction (F_(15,645)_ = 2.10, *p*=0.008, η_p_^2^ = 0.05, op = 0.97) but no main effect of group (F_(1,43)_ = 0.16, *p*=0.69, η_p_^2^ = 0.004, op = 0.07). Dividing the 16 sessions into 2, 8-session blocks; ANOVA found a block × group interaction (F_(1,43)_ = 4.81, *p*=0.034, η_p_^2^ = 0.10, op = 0.57). While sham (t_20_ = 7.69, p=2.13 × 10^−7^) and NAcc rats (t_23_ = 5.63, p=1.00 × 10^−5^) both increased poking from the first to second half of discrimination, sham rats showed greater increases (Figure 1E). Mean ± SEM baseline nose pokes rates for sessions 1-8: sham (28.44 ± 2.96) and NAcc (28.83 ± 1.97); sessions 9-16: sham (38.80 ± 3.62) and NAcc (35.33 ± 2.46; Figure 1E).

#### Fear Scaling

Sham rats acquired appropriate scaling of the fear response over the 16 sessions (Figure 2A, left). Suppression ratios for the entire 10-s cue were low in pre-exposure and initially increased to all cues. As discrimination proceeded, the suppression ratio for each cue diverged: high to danger, intermediate to uncertainty, and low to safety. NAcc rats showed a similar progression, but poorer overall scaling (Figure 2A, right). In support of the general emergence of scaling, ANOVA [between factor: group (sham vs. NAcc); within factors: session (16) and cue (danger, uncertainty and safety)] revealed a main effect of cue (F_(2,86)_ = 115.51, *p*=4.34 × 10^−25^, η_p_^2^ = 0.73, op = 1.00) and a cue × session interaction (F_(30,1290)_ = 14.05, *p*=6.30 × 10^−60^, η_p_ ^2^ = 0.25, op = 1.00). Revealing impaired scaling in NAcc rats, ANOVA found a cue × group interaction (F_(2,86)_ = 5.76, *p*=0.004, η_p_^2^ = 0.12, op = 0.86). The cue × group interaction was also observed when only the last six sessions were analyzed (F_(2,86)_ = 4.50, *p*=0.014, η_p_^2^ = 0.10, op = 0.76), the period by which scaling patterns were stable.

**Figure 2.**
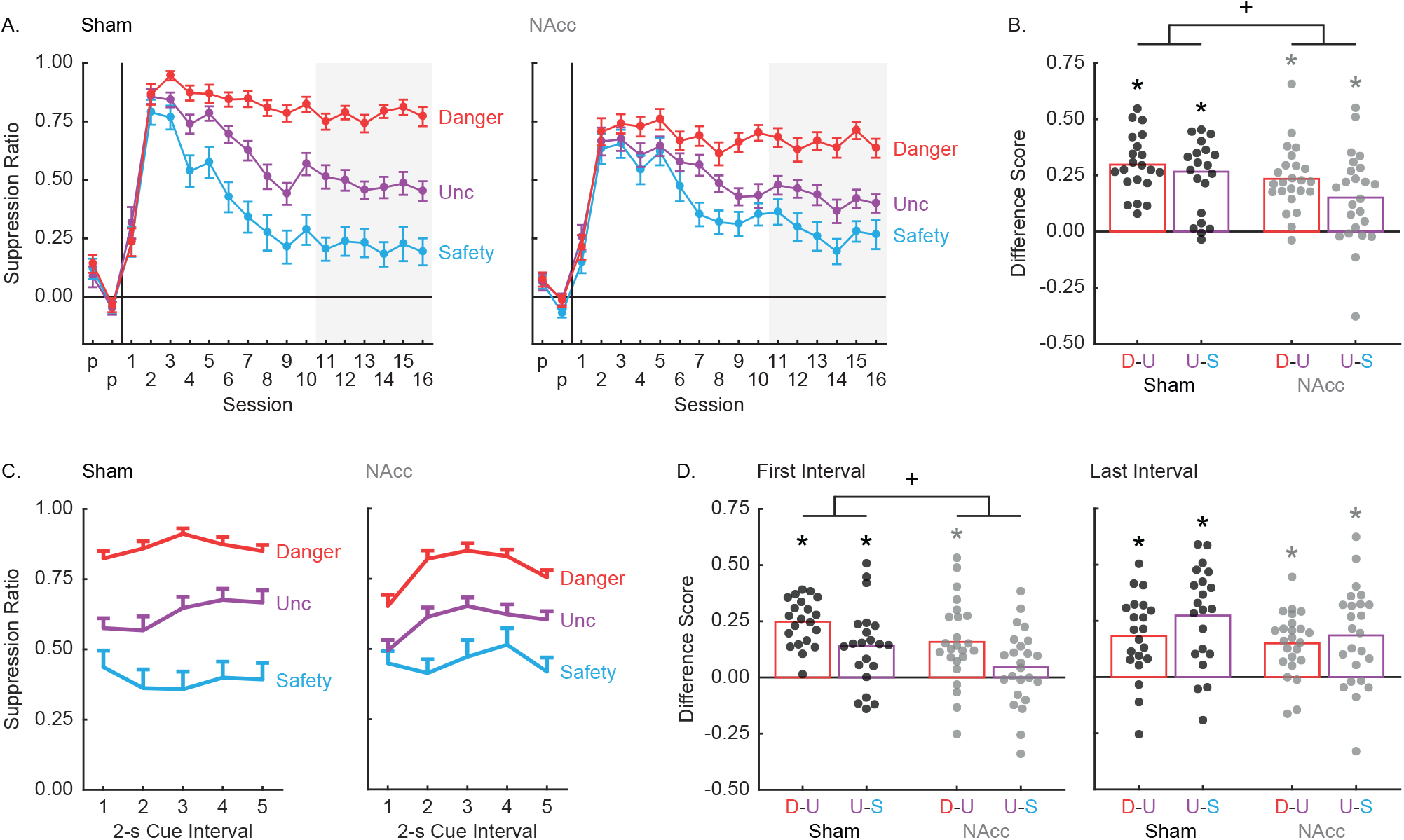
NAcc lesions and fear scaling. (A) Mean ± SEM suppression ratio for danger (red), uncertainty (purple), and safety (blue) are shown for sham (left) and NAcc (right) rats. The vertical lines separate the two pre-exposure and sixteen fear discrimination sessions. The last six discrimination sessions are shaded. (B) Mean difference score for danger vs. uncertainty (D-U, red bar) and uncertainty vs. safety (U-S, purple bar) across the entire 10-s cue is shown for sham (left) and NAcc (right) rats. Data points show individual difference scores. *One-sample t-test compared to zero, p<0.0125; +main effect of group, p<0.05. (C) Mean + SEM suppression ratios for the 5, 2-s cue intervals are shown for sham (left) and NAcc (right) rats. Cue color scheme maintained from A. (D) Mean difference score for danger vs. uncertainty (D-U, red bar) and uncertainty vs. safety (U-S, purple bar) is shown for the first 2-s cue interval (left) and last 2-s cue interval (right) for sham and NAcc rats. Data points show individual difference scores. *One-sample t-test compared to zero, p<0.00625; +main effect of group, p<0.05.

To further reveal the deficit in NAcc rats, we focused on suppression ratios from the final six sessions. Difference scores were calculated for the two components of scaling: (danger – uncertainty) and (uncertainty – safety). Sham (Figure 2B, left) and NAcc rats (Figure 2B, right) discriminated each cue pair. One-sample t-tests found that difference scores exceeded zero for each comparison: sham, danger vs. uncertainty (t_20_ = 10.25, *p*=2.07 × 10^−9^), uncertainty vs. safety (t_20_= 6.11, *p*=4.17 × 10^−8^); NAcc, danger vs. uncertainty (t_23_ = 8.01, *p*=0.001), uncertainty vs. safety (t_23_ = 3.65, *p*=0.002). However, NAcc rats showed poorer overall discrimination. ANOVA [between factor: group (sham vs. NAcc); within factor: discrimination (danger – uncertainty) and (uncertainty – safety)] revealed a main effect of group (F_(1,43)_ = 5.68, *p*=0.022, η_p_^2^ = 0.12, op = 0.64). Difference scores were reduced across both components in NAcc rats. These results reveal a general role for the NAcc in fear scaling.

#### Rapid Fear Scaling

We were interested in revealing a possible role for the NAcc in the rapid emergence of fear scaling. To do this, we examined mean suppression ratios from the last six sessions. Each cue was divided into 5, 2-s cue intervals and suppression ratios were calculated for each cue/interval. Sham rats showed scaling of the fear response in the first 2-s cue interval and in all subsequent intervals (Figure 2C, left). Scaling was reduced across all 2-s cue intervals in NAcc rats (Figure 2C, right). ANOVA [between factor: group (sham vs. NAcc); within factors: interval (5, 2-s cue intervals) and cue (danger, uncertainty and safety)] found a group × cue interaction (F_(2,86)_ = 3.88, *p*=0.024, η_p_^2^ = 0.08, op = 0.69). Supporting a specific role for the NAcc in rapid fear scaling, NAcc rats showed impaired scaling even when only the first 2-s cue interval was analyzed (cue × group interaction; F_(2,86)_ = 5.08, *p*=0.0008, η_p_^2^ = 0.11, op = 0.81). No cue × group interaction was observed when the last 2-s cue interval was analyzed (F_(2,86)_ = 1.90, *p*=0.16, η_p_^2^ = 0.04, op = 0.39).

To specify the nature of the deficit in NAcc rats, we reduced scaling into its component parts: (danger – uncertainty) and (uncertainty – safety). We calculated difference scores for the first and last 2-s cue intervals. Sham rats showed positive difference scores for each cue pair at each interval (Figure 2D, left). Difference scores exceeded zero, as revealed by one-sample t-tests: first 2-s cue interval: danger vs. uncertainty (t_20_ = 10.95, *p*=6.7 × 10^−4^), uncertainty vs. safety (t_20_ = 3.55, *p*=0.002); last 2-s cue interval: danger vs. uncertainty (t_23_ = 4.60, *p*=1.76 × 10^−4^), uncertainty vs. safety (t_23_ = 5.73, *p*=1.30 × 10^−5^) for shams. NAcc rats were generally impaired at rapid scaling. ANOVA for the first 2-s cue interval differences revealed a main effect of group (F_(1,43)_ = 6.50, *p*=0.014, η_p_^2^ = 0.01, op = 0.70), while ANOVA for the last 2-s cue interval differences scores found no main effect (F_(1,43)_ = 2.49, *p*=0.12, η_p_^2^ = 0.05, op = 0.34). Difference scores also suggest that NAcc rats were more specifically impaired in rapid uncertainty-safety discrimination (Figure 2D, right). One-sample tests found that only the NAcc uncertainty-safety difference score from the first 2-s cue interval failed to differ from zero: first interval: danger vs. uncertainty (t_23_ = 4.20, *p*=3.38 × 10^−4^), uncertainty vs. safety (t_23_ = 1.31, *p*=0.20); last interval: danger vs. uncertainty (t_20_ = 5.22, *p*=2.70 × 10^−5^), uncertainty vs. safety (t_20_ = 4.19, *p*=3.53 × 10^−4^). All significant, one-sample t tests survive Bonferroni correction (0.05/8, *p*<0.00625). Altogether, these results reveal a general role for the NAcc in the acquisition of rapid and overall fear scaling, as well as a more specific role in rapid uncertainty-safety discrimination.

### Experiment 2

Experiment 1 results implicate the NAcc in the acquisition of fear scaling. However, neurotoxic lesions altered baseline nose poking behavior and permanently ablated NAcc neurons. To determine a temporally specific role for the NAcc in expression, Experiment 2 took a within-subjects, optogenetic approach. Rats were NAcc-transducted with halorhodopsin or a control fluorophore, recovered, then acquired a scaled fear response to danger, uncertainty and safety. Once scaling was established, rats received sessions in which the NAcc was illuminated during cue presentation or during the inter-trial interval. If the NAcc plays identical roles in the acquisition and expression of fear scaling, we would expect to observe a three-way interaction (group × illumination × cue) with only halorhodopsin rats showing impaired overall scaling during cue illumination sessions. If the NAcc plays a more selective role in the expression of rapid fear scaling, we would anticipate a four-way interaction (group × interval × illumination × cue) with only halorhodopsin rats showing impaired rapid uncertainty-safety discrimination during cue illumination sessions.

#### Histological Results

Rats received bilateral NAcc transduction with halorhodopsin (Halo) or a control fluorophore (YFP) and bilateral optical ferrule implantation just above the NAcc. Representative transduction is shown (Figure 3A). Each subject’s total transduction area was drawn, made transparent, and stacked (Figure 3B). Darker areas indicate regions of greater overlap and more consistent transduction. Transduction centered around and above the anterior commissure, the precise NAcc location.

**Figure 3.**
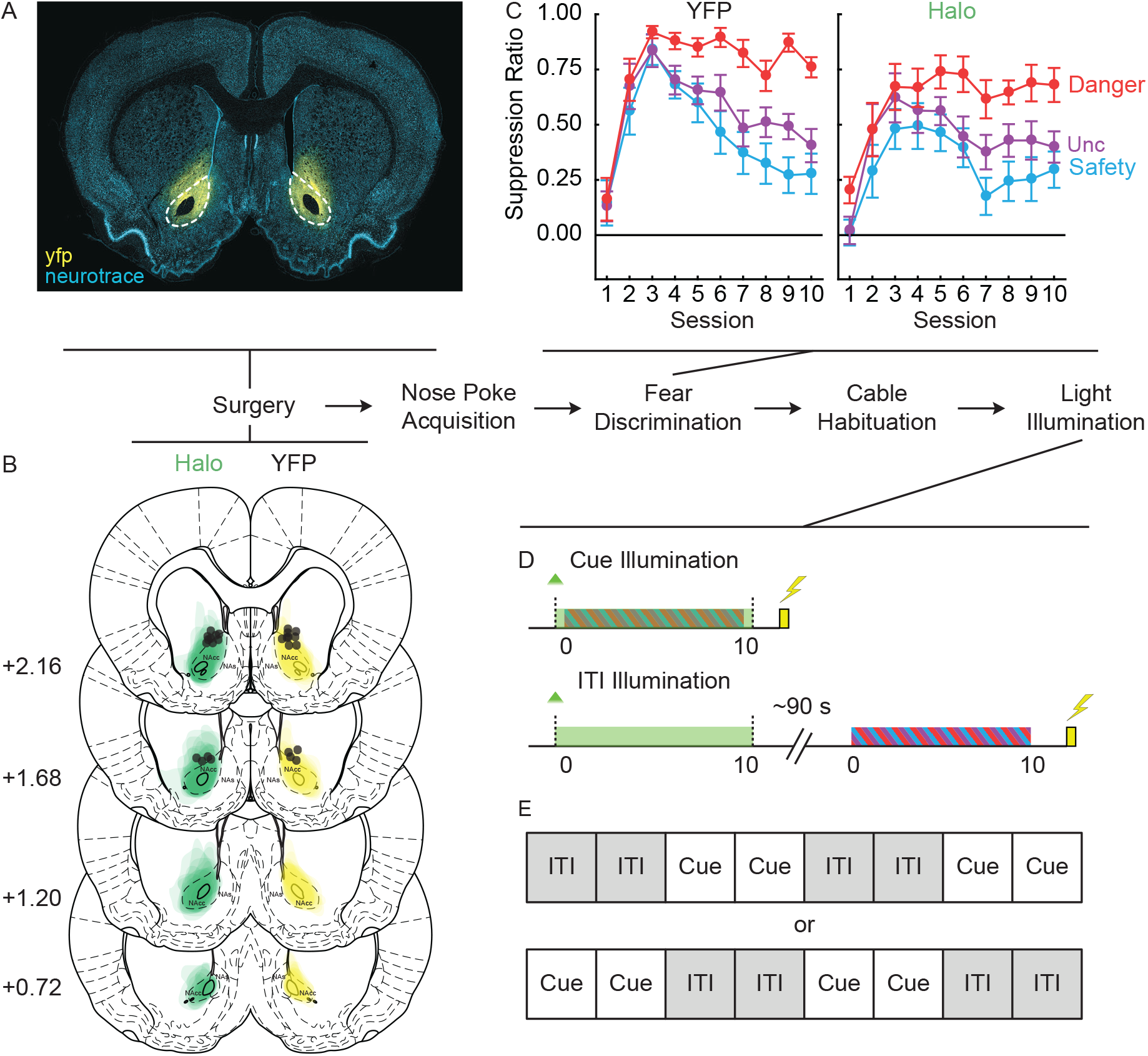
NAcc illumination experimental outline. (A) Representative NAcc transduction is shown with YFP expression (yellow fluorescent protein; yellow) and neurotrace (blue). Dotted lines indicate approximate NAcc location. (B) The extent of viral transduction across four coronal planes is shown for Halo (green, left) and YFP rats (yellow, right), and the anterior distance from bregma (millimeters) indicated. Individual ferrule placement is indicated in black circles. (C) Mean ± SEM suppression ratios for danger (red), uncertainty (purple), and safety (blue) are shown for YFP (left) and Halo rats (right) during the ten initial fear discrimination sessions. (D) In the final eight sessions, rats received NAcc light illumination during cue presentation (top) or during the inter-trial interval (ITI, bottom). Green indicates light illumination, yellow indicates shock delivery and candy-striped indicates cue presentation. (E) Cue and ITI illumination were given in alternating, two-session blocks. Block order was counterbalanced with roughly half of the subjects first receiving ITI illumination. Abbreviations: NAcc – nucleus accumbens core, NAs – nucleus accumbens shell.

#### Baseline Nose Poking

YFP and Halo rats showed equivalent baseline nose poking rates throughout pre-exposure, discrimination, cable habituation, and light illumination (Figure S1). ANOVA for baseline nose poke rate [factors: session (20) and group (YFP vs. Halo)] demonstrated a main effect of session (F_(19,437)_ = 12.60, *p*=4.19 × 10^−31^, η_p_^2^ = 0.35, op = 1.00), but no main effect or interaction with group (Fs < 0.93, *p*s>0.55). Equivalent performance lessens the concern that differences in suppression ratios between groups result from differences in baseline nose poke rates.

#### Initial Fear Scaling

YFP and Halo rats acquired reliable fear scaling over the 10 sessions (Figure 3C). Suppression ratios were low in pre-exposure and initially increased to all cues. As discrimination proceeded, the suppression ratio for each cue diverged: high to danger, intermediate to uncertainty, and low to safety. Demonstrating overall scaling, ANOVA [within factors: session (10) and 10-s cue (danger, uncertainty and safety); between factor: group (YFP vs. Halo)] revealed a main effect of cue (F_(2.46)_ = 36.21, *p*=3.58 × 10^−10^, η_p_^2^ = 0.61, op = 1.00), session (F_(9,207)_ = 25.74, *p*=2.04 × 10^−29^ η_p_^2^ = 0.53, op = 1.00) and a cue × session interaction (F_(18,414)_ = 6.26, *p*=1.14 × 10^−13^, η_p_^2^ = 0.21, op = 1.00). ANOVA found no main effect or interaction with group (Fs < 3.42, *p*s>0.08). Thus, YFP and Halo entered the light illumination phase (Figure 3D, E) showing equivalent fear scaling.

#### Overall Fear Scaling during Light Illumination

When suppression ratios were calculated for the entire 10-s cue, YFP and Halo rats showed scaling of the fear response over the 10 sessions of cable habituation, cue illumination and ITI illumination (Figure 4). ANOVA [between factor: group (YFP vs. Halo); within factors: session (10) and cue (danger, uncertainty and safety)] was separately performed for rats receiving ITI-cue illumination order (YFP, n = 5; Halo, n = 5; Figure 4A) and cue-ITI illumination order (YFP, n = 7; Halo, n = 8; Figure 4B). Each ANOVA returned a main effect of cue (Fs > 29, *ps*<2 × 10^−7^), but neither returned a main effect of group, group × cue interaction or a group × cue × session interaction (Fs < 2.5, ps>0.1). Complete ANOVA results provided in Table 1. Next we calculated difference scores for the two components of scaling: (danger – uncertainty) and (uncertainty – safety) (Figure 4C). ANOVA [between factors: group (YFP vs. Halo) and order (ITI-cue vs. cue-ITI); within factors: illumination (hab/ITI vs. cue) and discrimination (danger – uncertainty vs. uncertainty – safety)] found main effects of illumination (F_(1,21)_ = 8.90, *p*=0.007, η_p_^2^ = 0.30, op = 0.81) and discrimination (F_(1,21)_ = 14.29, *p*=0.001, η_p_^2^ = 0.41, op = 0.95), as well as a group × illumination interaction (F_(1,21)_ = 4.75, *p*=0.041, η_p_^2^ = 0.19, op = 0.55). The interaction resulted from YFP rats showing poorer overall discrimination in cue illumination sessions compared to ITI illumination, whereas Halo rats showed equivalent discrimination in each session type. No main effect of group (F_(1,21)_ = 0.19, *p*=0.67, η_p_^2^ = 0.009, op = 0.07) or any group interaction was detected (Fs < 1.2, ps>0.3). These results reveal that NAcc activity is not necessary for the expression of fear scaling when suppression is measured for the duration of cues.

**Table 1.** ANOVA results for 10-s cue.

| Term | F | p | $\eta_p^2$ | op |
| --- | --- | --- | --- | --- |
| **cue** | **103.37** | **$7.09 \times 10^{-10}$** | **0.93** | **1.00** |
| cue x group | 0.55 | 0.59 | 0.06 | 0.13 |
| session | 1.55 | 0.15 | 0.16 | 0.68 |
| **session x group** | **2.04** | **0.047** | **0.2** | **0.82** |
| cue x session | 1.21 | 0.26 | 0.13 | 0.79 |
| cue x session x group | 1.18 | 0.29 | 0.13 | 0.78 |
| group | 2.51 | 0.15 | 0.24 | 0.29 |

10-s cue ANOVA results Cue-ITI
| Term | F | p | $\eta_p^2$ | op |
| --- | --- | --- | --- | --- |
| **cue** | **29.23** | **$2.23 \times 10^{-7}$** | **0.69** | **1.00** |
| cue x group | 0.15 | 0.86 | 0.01 | 0.07 |
| **session** | **3.46** | **0.001** | **0.21** | **0.98** |
| session x group | 0.77 | 0.65 | 0.06 | 0.37 |
| cue x session | 1.07 | 0.39 | 0.08 | 0.74 |
| cue x session x group | 1.47 | 0.10 | 0.10 | 0.90 |
| group | 1.19 | 0.30 | 0.08 | 0.17 |

**Figure 4.**
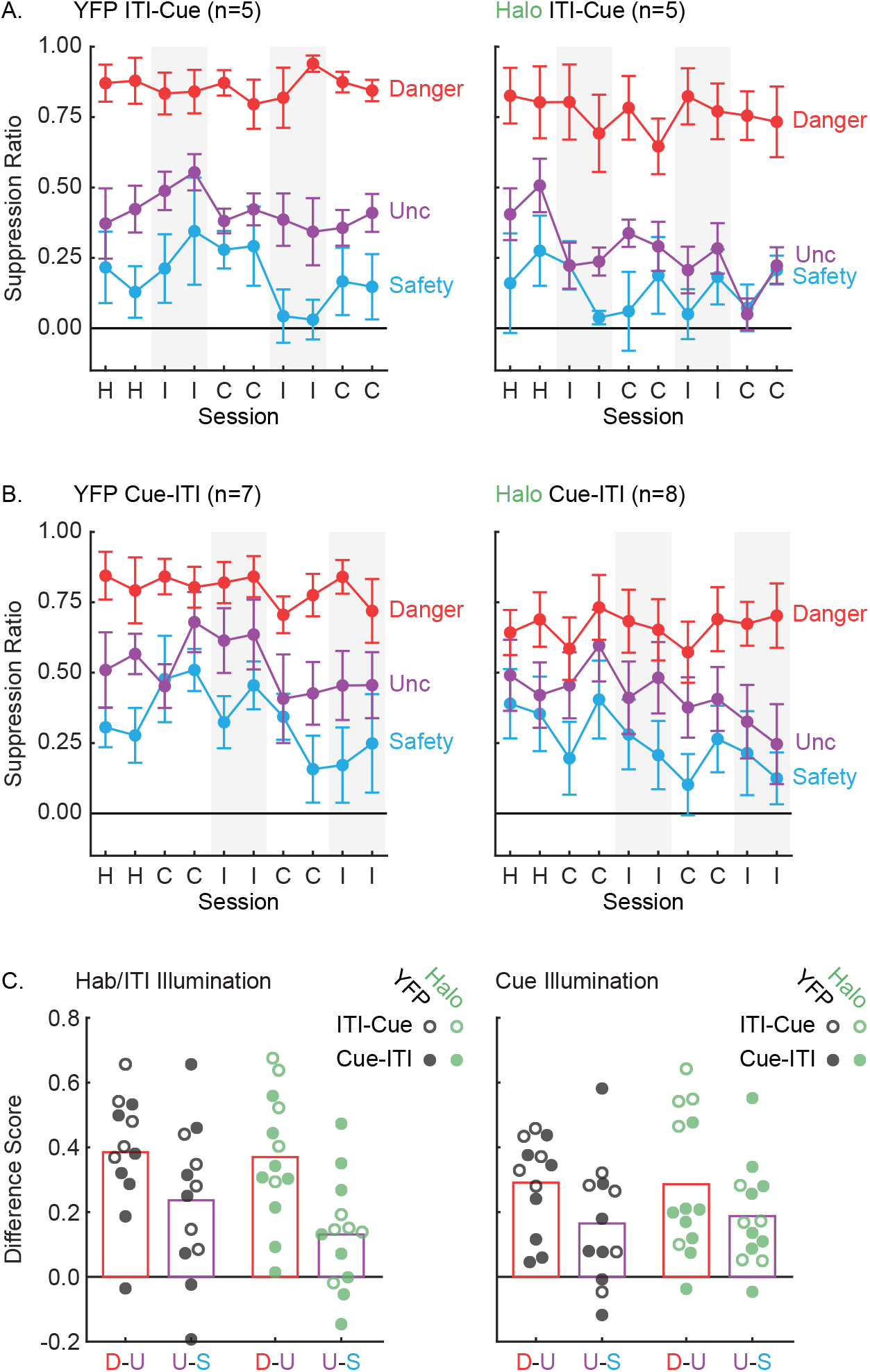
NAcc illumination and overall fear scaling. (A) Mean ± SEM suppression ratios over the entire 10-s cue are plotted for danger (red), uncertainty (purple), and safety (blue). Data are plotted for cable habituation (H), ITI illumination (I) and cue illumination (C) for YFP (n=5) and Halo rats (n=5) receiving ITI-cue illumination. ITI illumination sessions shaded. (B) YFP (n=7) and Halo rats (n=8) receiving cue-ITI illumination plotted as in A. (C) Difference scores for danger vs. uncertainty (D-U, red bar) and uncertainty vs. safety (U-S, purple bar) are shown for YFP (black) and Halo rats (green) during cable habituation/ITI illumination (left) and cue illumination (right). ITI-cue rats indicated by open circles, cue-ITI rats by closed circles.

#### Rapid Fear Scaling during Light Illumination

To examine rapid fear scaling, we divided the 10-s cue into 5, 2-s intervals. Suppression ratios are shown for each cue/interval during habituation/ITI illumination sessions [YFP rats (Figure 5A) and Halo rats (Figure 5B)] and for cue illumination sessions [YFP rats (Figure 5C) and Halo rats (Figure 5D)]. To examine a possible role for the NAcc in rapid fear scaling, we performed ANOVA with all factors [within factors: session-type (cable habituation, ITI illumination and cue illumination), cue (danger, uncertainty and safety), and interval (5, 2-s cue intervals); between factor: group (YFP vs. Halo)]. The complete ANOVA output is reported in Table 2. Consistent with general scaling across groups, ANOVA revealed a main effect of cue (F_(2,46)_ = 89.04, *p*=1.53 × 10^−16^, η_p_^2^ = 0.80, op = 1.00) as well as a cue × interval interaction (F_(8,184)_ = 6.14, *p*=5.16 × 10^−7^, η_p_^2^ = 0.21, op = 1.00). Indicative of a selective role for the NAcc in rapid fear scaling, ANOVA revealed a significant 4-way interaction [session-type × cue × interval × group (F_(16,368)_ = 1.80, *p*=0.029, η_p_^2^ = 0.07, op = 0.95)], but not a significant 3-way interaction [session-type × cue × group (F_(4,92)_ = 1.35, *p*=0.26, η_p_^2^ = 0.06, op = 0.41)].

**Table 2.** ANOVA results for 5, 2-s cue intervals.

| Term | F | p | $\eta_p^2$ | op |
| --- | --- | --- | --- | --- |
| group | 1.94 | 0.18 | 0.08 | 0.27 |
| **session-type** | **10.64** | **1.59 x 10-4** | **0.32** | **0.99** |
| session-type x group | 2.22 | 0.12 | 0.09 | 0.43 |
| **cue** | **89.04** | **1.53 x 10-16** | **0.80** | **1.00** |
| cue x group | 0.57 | 0.57 | 0.02 | 0.14 |
| **interval** | **6.00** | **2.46 x 10-4** | **0.21** | **0.98** |
| interval x group | 1.37 | 0.25 | 0.06 | 0.41 |
| **session-type x cue** | **4.21** | **0.004** | **0.16** | **0.91** |
| session-type x cue x group | 1.35 | 0.26 | 0.06 | 0.41 |
| session-type x interval | 0.71 | 0.68 | 0.03 | 0.32 |
| session-type x interval x group | 1.46 | 0.17 | 0.06 | 0.65 |
| **cue x interval** | **6.14** | **5.16 x 10-7** | **0.21** | **1.00** |
| cue x interval x group | 0.64 | 0.74 | 0.03 | 0.29 |
| session-type x cue x interval | 0.78 | 0.71 | 0.03 | 0.54 |
| **session-type x cue x interval x group** | **1.80** | **0.029** | **0.07** | **0.95** |

**Figure 5.**
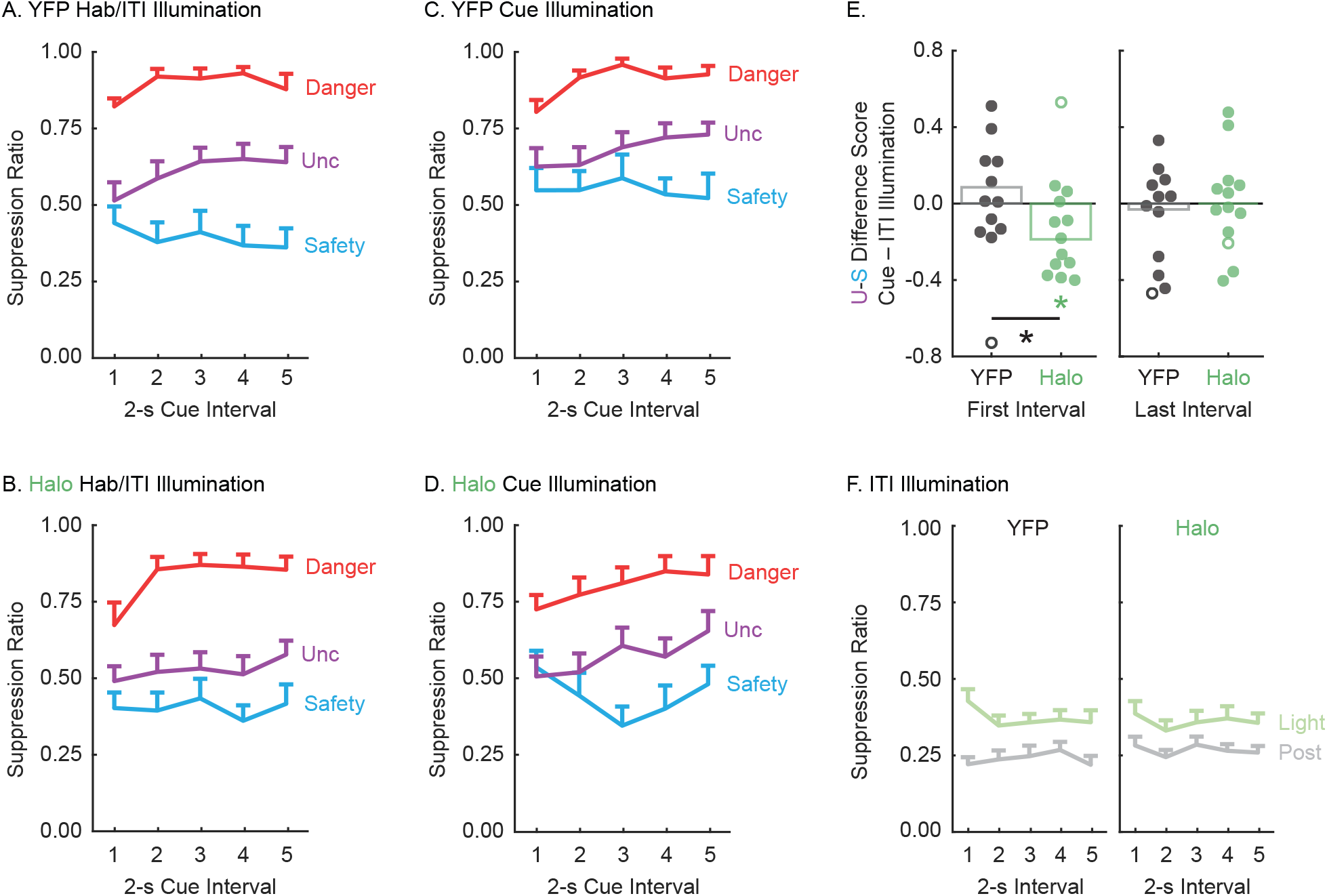
NAcc illumination and rapid fear scaling. Mean + SEM suppression ratios are plotted for the 5, 2-s cue intervals for danger (red), uncertainty (purple), and safety (blue), for YFP and Halo rats during (A & B) cable habituation/ITI illumination, and (C & D) cue illumination. (E) Uncertainty – safety difference scores were separately calculated for cue and ITI illumination, then difference of difference scores calculated (cue difference score – ITI difference score). Mean and individual difference of difference scores are plotted for the first 2-s cue interval (left) and last 2-s cue interval (right), for YFP (black) and Halo rats (green). Open circles are outliers. *(green) one-sample t-test compared to zero, p=0.0038. *(black) independent samples t-test, p=0.0041. (F) Mean + SEM suppression ratios are plotted for the 5, 2-s intervals during ITI illumination (light green) and for the 5, 2-s intervals during the post-illumination period (gray) (YFP, left; Halo, right).

The 4-way interaction indicates that YFP and Halo rats showed differing temporal scaling patterns across the different session types. To begin to clarify the differing patterns, we split YFP and Halo rats and performed identical ANOVAs [within factors: session-type (habituation, ITI illumination and cue illumination), cue (danger, uncertainty and safety), and interval (5, 2-s cue intervals)]. Indicative of reliable scaling, ANOVA for YFP rats found a main effect of cue (F_(2,22)_ = 47.71, *p*=1.0 × 10^−10^, η_p_^2^ = 0.81, op = 1.00) and a cue × interval interaction (F_(8,88)_ = 2.76, *p*=0.009, η_p_ ^2^ = 0.20, op = 0.92). Revealing no effect of illumination on the temporal pattern of fear scaling, the 3-way interaction (session-type × cue × interval) was not significant (F_(16,176)_ = 0.59, *p* = 0.89, η_p_^2^ = 0.05, op = 0.39). ANOVA for Halo rats also found a main effect of cue (F_(2,24)_ = 41.39, *p*=1.66 × 10^−8^, η_p_^2^ = 0.78, op = 1.00) and a cue × interval interaction (F_(8,96)_ = 4.07, *p*=3.36 × 10^−4^, η_p_^2^ = 0.25, op = 0.99). Only now, ANOVA revealed a significant 3-way interaction (session-type × cue × interval; F_(16,92)_ = 1.92, *p* = 0.021, η_p_^2^ = 0.14, op = 0.95). NAcc illumination only disrupted the temporal scaling pattern for Halo rats. It appears that – similar to NAcc rats – Halo rats receiving NAcc optogenetic inhibition during cue presentation were specifically impaired in rapid uncertainty-safety discrimination (Figure 5D). If this were the case, then Halo rats should show poorer uncertainty-safety discrimination in the first 2-s cue interval during cue illumination sessions compared to ITI illumination sessions. YFP rats show would equivalent performance during each type of illumination. Further, this deficit should not be observed in the last 2-s cue interval.

We calculated (uncertainty – safety) difference scores for the first and last 2-s cue intervals. Separate scores were calculated for cue and ITI illumination sessions. We then calculated a difference score for the two session-illumination-types (cue difference score – ITI difference score). This approach capitalized on our within-subject design; each rat was tested during cue and ITI illumination. The approach is consistent with our ANOVA results, which found a differential effect of cue and ITI illumination for Halo rats, but not for YFP rats. A difference score of difference scores has the added benefit of reducing the differential illumination effects to a single value. Values around zero would indicate equivalent uncertainty-safety discrimination during cue and ITI illumination sessions. Negative values would indicate *worse* uncertainty-safety discrimination during cue illumination sessions. Two individuals (1 YFP and 1 Halo) had first interval difference scores ±2 standard deviations beyond the group mean. The data for these individuals are shown (Figure 5E, open circles), but were not included in t-test analyses.

In the first 2-s cue interval, Halo rats showed worse uncertainty-safety discrimination during cue illumination sessions compared to ITI illumination sessions (Figure 5E, left). This was supported by significant, negative shift of differences scores away from zero (one-sample t-test, t_11_ = −3.65, *p*=0.004). YFP rats showed equivalent uncertainty-safety discrimination during cue and ITI illumination sessions; difference scores hovered around zero (t_10_ = 1.22, *p*=0.25). Further, YFP and Halo difference scores differed from one another (independent samples t-test, t_21_ = 3.22, p=0.004). Impaired uncertainty-safety discrimination in Halo rats receiving cue illumination was restricted to the first 2-s cue interval. Identical analysis of the last 2-s cue interval found that difference scores did not differ from zero for YFP (one-sample t-test, t_10_ = −0.41, *p*=0.69) and Halo rats (one-sample t-test, t_11_ = 0.27, *p*=0.80) (Figure 5E, right). Differences scores were similar between the two groups (independent samples t-test, t_21_ = 0.48, p=0.64). Altogether, the results reveal that NAcc activity at the time of cue presentation is necessary to rapidly discriminate uncertainty and safety.

Of course, it is possible that NAcc optogenetic inhibition simply suppressed rewarded nose poking. In this case, impaired rapid fear scaling would be the byproduct of a general reduction in poking. To rule out this possibility, we examined nose poke suppression during light illumination in ITI sessions (Figure 5F). No cues were present during this period, allowing us to determine the effect of light illumination alone to suppress nose poking. The middle 10 s of the 11-s light illumination was divided into 5, 2-s cue intervals – exactly as was done for the cue illumination analyses. For comparison, we also sampled 10 s of nose poking 30-s following illumination offset. This post-illumination served as a control period to which light illumination could be compared. ANOVA [within factors: period (light and post) and interval (5, 2-s cue intervals); between factor: group (YFP vs. Halo)] revealed main effects of period (F_(1.23)_ = 34.53, *p*=5 × 10^−6^, η_p_^2^ = 0.60, op = 1.00) and interval (F_(4,92)_ = 2.49, *p*=0.049, η_p_^2^ = 0.10, op = 0.69). Critically, ANOVA found no main effect or interaction with group (Fs < 1.10, *p*s>0.31). So while suppression ratios were higher during light illumination, this did not differ between YFP and Halo rats and was therefore not due to inhibition of NAcc activity.

## Discussion

We set out to examine a role for the nucleus accumbens core in fear scaling. Neurotoxic lesions revealed a general role for the NAcc in the acquisition of fear scaling, as well as a specific role in acquiring rapid uncertainty-safety discrimination. Optogenetic inhibition revealed a specific role for NAcc cue activity in the expression of rapid, uncertainty-safety discrimination. The results reveal that the NAcc is an essential component of a neural circuit permitting fear to rapidly scale to degree of threat.

Before discussing our results more broadly, we must consider several limitations of our experimental design and results. First, our experiments only used male rats. Several studies have reported sex differences in danger-safety discrimination (Day et al., 2016; Foilb et al., 2018; Greiner et al., 2019). We find only modest sex differences in our discrimination procedure (Walker et al., 2018; Walker et al., 2019), suggesting similar neural circuits may be utilized across sexes. Of course, females and males may achieve similar performance through differing neural mechanisms. Another important consideration is that our dependent measure of fear is derived from the rate of rewarded nose poking. Conditioned suppression is a strength because it provides an objective measure of fear on multiple time scales (Estes and Skinner, 1941; Bouton and Bolles, 1980). It is a potential weakness because the NAcc plays a well-established role in reward-seeking. Disrupting NAcc function can attenuate reward-related behavior in many settings (Corbit et al., 2001; Hall et al., 2001; Ito et al., 2004; Blaiss and Janak, 2009; Ambroggi et al., 2011; McDannald et al., 2011; McDannald et al., 2013), though this finding is not universal (Ramirez and Savage, 2007; Corbit and Balleine, 2011). In our first experiment, NAcc lesions slowed the increase of baseline nose poking over discrimination sessions and also impaired fear scaling. However, the temporal emergence of the deficits in reward-seeking and fear scaling did not align. The fear scaling deficit was apparent across all sessions while the nose poking deficit only emerged in the later sessions.

Our second experiment better demonstrated independent roles for the NAcc in rapid fear scaling and reward-seeking. Light illumination during the cue period impaired rapid uncertainty-safety discrimination in NAcc-halorhodopsin rats, but not NAcc-YFP rats. By contrast, light illumination during the inter-trial interval produced equivalent and modest reductions in nose poking for both groups. Optogenetic inhibition of the NAcc was insufficient to reduce rewarded nose poking. The failure of NAcc inhibition to suppress nose poking may seem odd. Mice will readily perform actions that channelrhodopsin-excite NAcc D1 and D2 cell types (Cole et al., 2018), and rats will perform actions exciting glutamatergic inputs to the NAcc via channelrhodopsin (Stuber et al., 2011; Britt et al., 2012). However, these studies demonstrate that NAcc activity is sufficient, but not necessary, to support reward-seeking. Further, prominent theories posit that reward-seeking initially depends on medial striatal structures, such as the NAcc. With further training, lateral striatal regions, such as the dorsolateral striatum, control reward-seeking (Gerdeman et al., 2003; Belin and Everitt, 2008; Corbit et al., 2012; Burton et al., 2015). In our second experiment, rats had extensive experience with nose poking by the time the NAcc was optogenetically inhibited. By this time, reward-seeking may have no longer been under NAcc control, yet the NAcc continued to contribute to rapid fear scaling.

A role for the NAcc in fear would be expected based on immediate early gene studies. Shock-associated cues and contexts reliably upregulate NAcc c-fos and zif268 (Beck and Fibiger, 1995; Campeau et al., 1997; Thomas et al., 2002). Despite these clear findings, specifying the role of the NAcc in fear has presented a considerable challenge. Initial work by Parkinson and colleagues found that NAcc lesions impaired cued fear, but enhanced contextual fear (Parkinson et al., 1999). Taking a similar experimental approach, Levita and colleagues found that NAcc lesions had no impact on the acquisition or expression of cued fear, but impaired retention of contextual fear (Levita et al., 2002). Contemporary work by Haralambous and Westbrook found that inhibiting accumbens activity (core + shell) specifically impaired the acquisition, but not expression of contextual fear, and had no effect on cued fear (Haralambous and Westbrook, 1999). Even considering slightly different methodologies, it is difficult to reconcile these disparate results.

These are not the only conflicts in the literature. Schwienbacher and colleagues found that blocking NAcc activity with tetrodotoxin abolished acquisition, and impaired expression, of fear-potentiated startle (Schwienbacher et al., 2004). The very next year, Josselyn and colleagues utilized a variety of methods to manipulate the NAcc during fear-potentiated startle: lesion, agonizing dopamine and blocking glutamate. NAcc manipulation had no effect on any aspect of fear-potentiated startle (Josselyn et al., 2005). Since these initial studies, the NAcc has been implicated in a variety of fear-related processes. For example, the NAcc can modulate salience, regulating the ability of cues to enter into associations with shock (Iordanova et al., 2006a; Iordanova et al., 2006b; Iordanova, 2009). Human imaging studies have observed correlates of prediction error, a theoretical signal that strengthens or weakens cue-shock associations (Seymour et al., 2004; Delgado et al., 2008; Schiller et al., 2008; Li et al., 2011), consistent with a role for the NAcc in predictive learning (Li and McNally, 2015).

What can we make of the mixed NAcc fear literature? Though dissatisfying, one answer is that the NAcc must play multiple roles in fear. Genetically- and anatomically-defined NAcc neuron types may be linked to specific fear processes. Future work dissecting the NAcc in this way, as has been done in reward settings (Kupchik et al., 2015; Francis and Lobo, 2017; Tejeda et al., 2017), is likely to be fruitful. A more fulfilling answer might be that standard cued and contextual fear conditioning procedures do not isolate essential NAcc functions. The NAcc may not be needed to demonstrate fear to certain threat or to withhold fear to certain safety. For example, the NAcc is not necessary to behaviorally discriminate contexts/cues associated with certain shock and certain safety (Antoniadis and McDonald, 2006; McDannald and Galarce, 2011; Piantadosi, 2017).

We propose that a necessary role for the NAcc in fear emerges when subjects are confronted with threats on a continuum from safety to danger. The NAcc is a core component of a neural circuit permitting the level of the fear response to scale to the degree of threat. Further, our results suggest general and specific roles for the NAcc in fear scaling. During acquisition, the NAcc is generally necessary for fear scaling for the duration of an encounter – in our case for the entirety of cue presentation. At the same time, the NAcc is specifically necessary for one component of fear scaling: rapid discrimination of uncertain threat and safety. Once a scaled fear response is acquired, the general role for the NAcc diminishes. However, the NAcc continues to play a specific role in rapidly discriminating uncertain threat and safety. Of course, we are not claiming that fear scaling is *the* function of the NAcc in fear, but rather *a* function.

Environmental threats are not absolute, but exist on a continuum from safety to danger. Using a behavioral procedure that attempts to capture this continuum, we find the NAcc is essential to scale fear to degree of threat. Our results clarify at least one role for the NAcc in fear, yet much more work remains. There is evidence of altered NAcc structure and function in anxiety and stress disorders (Cha et al., 2014; Felmingham et al., 2014; Manning et al., 2015; Morey et al., 2017). Disrupted threat-safety discrimination may be conceptualized as maladaptive fear scaling. Detailing NAcc function, and mapping a more complete neural circuit for rapid fear scaling, may inform strategies to promote adaptive fear in anxiety and stress disorders.

## Acknowledgements

We would like to thank Alexa LaBanca and Andrew Thomson for technical support. This work was supported by NIH DA034010 to M.A.M. We thank Bret Judson and the Boston College Imaging Core for infrastructure and support.

---

**Figure S1.**
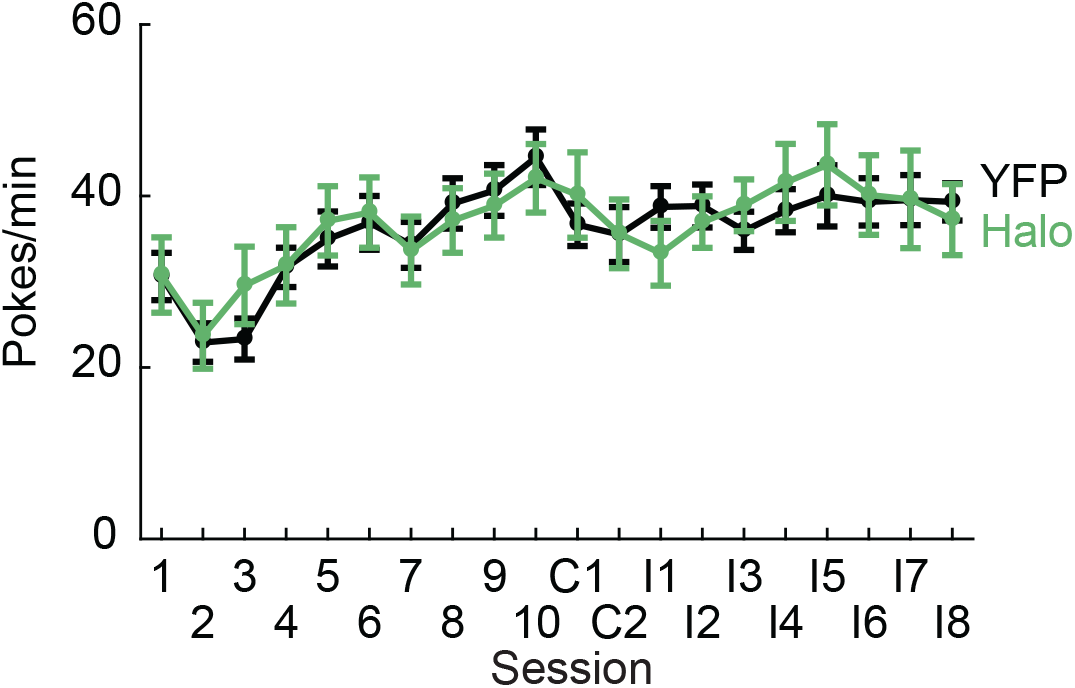
Baseline nose poking. Mean +/- SEM nose poke rate (y-axis) is shown YFP (black) and Halo rats (green) during the 10 pre-illumination, 2 cable habituation (C) and 8 illlumination (I) sessions. Repeated measures ANOVA (between factor: group; within factor: session) found neither a main effect of group nor a group × session interaction.

